# Neurotransmitter content heterogeneity within an interneuron class shapes inhibitory transmission at a central synapse

**DOI:** 10.1101/2022.05.03.490471

**Authors:** Dimitri Dumontier, Caroline Mailhes-Hamon, Stéphane Supplisson, Stéphane Dieudonné

## Abstract

Neurotransmitter content is deemed the most basic defining criterion for neuronal classes, contrasting with the intercellular heterogeneity of many other molecular and functional features. Here we show, in the adult mouse brain, that neurotransmitter content variegation within a neuronal class is a component of its functional heterogeneity. Most Golgi cells (GoCs), the well-defined class of cerebellar interneurons inhibiting granule cells (GrCs), contain cytosolic glycine, accumulated by the neuronal transporter GlyT2, and GABA in various proportions. To assess the functional consequence of this neurotransmitter variation, we paired GrCs recordings with optogenetic stimulations of single GoCs, which preserve the intracellular transmitter mixture. We show that the strength and decay kinetics of GrCs IPSCs, which are entirely mediated by GABAA receptors are negatively correlated to the presynaptic expression of GlyT2 by GoCs. We isolate a slow spillover component of GrCs inhibition that is also affected by the expression of GlyT2, leading to a 56 % decrease in relative charge. Acute manipulations of cytosolic GABA and glycine supply recapitulate the modulation of IPSC charge, supporting the hypothesis that presynaptic loading of glycine negatively impact the GABAergic transmission in mixed interneurons through a competition for vesicular filling. Our results suggest that heterogeneity of neurotransmitter supply within the GoC class may provide a presynaptic mechanism to tune the gain of the stereotypic granular layer microcircuit, thereby expanding the realm of possible dynamic behavior.

## Introduction

Cellular neuroscience was founded on the morphological identification of neuronal classes, based on their dendritic and axonal morphologies, as revealed by Golgi staining. Refining this classification using electrophysiological and molecular profiles and assigning functions to individual neuron classes is still a central effort of modern neuroscience. However, adding more parameters to the description of neurons deceptively resulted in conflicting class separations (Petilla Interneuron Nomenclature et al 2008, Tremblay et al 2016, Zeng & Sanes 2017), leading in the most extreme cases to a fractal view of neuronal populations (Hobert et al 2016, Parra et al 1998). Recently, single-cell RNA sequencing approaches have enabled clustering into broad neuronal classes, within which the large residual intra-class variability may not be easily reconciled with previously identified subpopulations (Kozareva et al 2021, Saunders et al 2018, Tasic et al 2016, Zeisel et al 2015).

Amidst this complexity, the neurotransmitter released by each neuron has been considered a reliable criterion for defining broad neuronal classes. It delineates excitatory, inhibitory and neuromodulatory neuronal populations and has been extended to neuropeptides, such as somatostatin and Vasointestinal peptide, to classify forebrain interneurons (Kepecs & Fishell 2014, Lim et al 2018, Tremblay et al 2016). However, the prevalence of neurons releasing multiple neurotransmitters or neuromodulators (Granger et al 2017) has somewhat blurred this simple view, opening the possibility for a morpho-functional neuronal class to contain cells with quantitatively or qualitatively different mixture of neurotransmitters, with or without immediate postsynaptic receptors. This constitutes a dazzling prospect, whereby the synaptic output of each presynaptic neuron within a class, defined presynaptically by its neurotransmitter mixture, could be coordinated with its other properties such as its excitability or sensitivity to neuromodulation.

Varied cytosolic accumulation and synaptic co-release of GABA and glycine, as occurs in the majority of hindbrain inhibitory neurons, constitutes the most studied case of variability in neurotransmitter content (Awatramani et al 2005, Chery & de Koninck 1999, Dugue et al 2005, Dumoulin et al 2001, Giber et al 2015, Jonas et al 1998, Moore & Trussell 2017, Nerlich et al 2017, O’Brien & Berger 1999). GABAergic, glycinergic and mixed inhibitory neurons persist into adulthood, when they can contact the same postsynaptic neurons (Batten et al 2010, Dufour et al 2010, Dumba et al 1998, Husson et al 2014, Lu et al 2008, Ottersen et al 1988, Paik et al 2019, Riquelme et al 2001, Todd & Sullivan 1990). However, to date, the diversity of IPSCs properties has always been attributed to the expression by the postsynaptic neurons of different flavors of GABA and glycine receptors (Ahmadi et al 2003, Awatramani et al 2005, Chery & de Koninck 1999, Chery & De Koninck 2000, Dugue et al 2005, Dumoulin et al 2001, Gao et al 2001, Giber et al 2015, Gonzalez-Forero & Alvarez 2005, Keller et al 2001, Lu et al 2008, Mitchell & Silver 2000, Moore & Trussell 2017, Nerlich et al 2017, Rossi & Hamann 1998, Rousseau et al 2012). While presynaptic and postsynaptic co-maturation has been reported (Nabekura et al 2004), the role of presynaptic inhibitory transmitter content in the properties and strength of mature synapses has been overlooked. It remains to be formally investigated whether inhibitory neurons of the same morpho-functional class use different GABA/glycine mixture and how this mixture alters the characteristics of their postsynaptic action.

To address this question, we studied a well-defined morphological class of interneurons in the cerebellar cortex, the Golgi cells (GoCs), which exhibit stereotyped morphology and connectivity (Dieudonne 1998, Galliano et al 2010) but diversity in neurotransmitter content (Dugue et al 2005, Ottersen et al 1987, Ottersen et al 1988). GoCs span the full spectrum with a fifth of pure GABAergic neurons, which express only the GABA synthesis enzyme (glutamate decarboxylase GAD), a majority of mixed GoCs that additionally express the neuronal transporter of glycine, GlyT2, and a handful of pure glycinergic cells that express GlyT2 but not GAD (Ottersen et al 1987, Simat et al 2007).

This diversity is also observed at axonal varicosities of GoCs in cerebellar glomeruli (Dugue et al 2005, Ottersen et al 1988), where they synapse on billions of granule cells (GrCs) and on rare unipolar brush cells (UBCs). Studies of inhibition between GoCs and UBCs (Dugue et al 2005, Rousseau et al 2012) have demonstrated a major role for differential expression of GABAARs and GlyRs by postsynaptic UBCs in shifting transmission from a mixed system to a pure GABAergic system, with no evidence of an impact of presynaptic neurochemistry (Rousseau et al 2012), as in other mixed systems to date. Remarkably, GrCs express only GABAAR, with two well-established subunit compositions: low-affinity synaptic receptors mediating fast IPSCs and high-affinity extrasynaptic receptors mediating a slow spillover component (Dumoulin et al 2001, Kaneda et al 1995, Rossi & Hamann 1998, Wall & Usowicz 1997). The absence of postsynaptic GlyR constitutes somewhat counterintuitively, by eliminating the dominant effect of postsynaptic receptor variability, an asset to explore the functional impact of presynaptic GABA/glycine content diversity.

In this paper we performed optogenetic stimulations of phenotypically-identified GoCs and paired whole-cell GrCs recordings. We show that marker of GlyT2 expression level in GoCs are inversely correlated with the synaptic strength (kinetics and amplitude) of the GrC IPSCs. We demonstrate, by manipulating the supply of GABA and glycine to GoCs, the existence of a presynaptic mechanism of synaptic strength modulation. Overall, we present evidence that variability in transmitter content within a neuronal class may be an important component of the functional variability of that class and discuss the implications of this finding.

## Results

### A targeted optogenetic stimulation strategy for GoC-GrC paired recordings without presynaptic dialysis

To investigate the impact of each GoCs neurotransmitter mixture on its functional output, we sought to record pairs between neurochemically-identified GoCs and their synaptically connected GrCs targets. Whole-cell pair recordings, which represent the technique of choice to characterize synaptic connections, cannot be used in this case, because dialysis of the cytoplasm of the presynaptic GoCs would alter its neurotransmitter content and synaptic physiology (Diana & Marty 2003, Wang et al 2013). To address this issue, we implemented targeted optogenetic stimulations of GoCs. Channelrhodopsin (ChR2) expression in identified GoCs was achieved by stereotactic injections of CRE-dependent AAV2.1 virus into the cerebellum of GlyT2-Cre or GlyT2-eGFP mice (**Figure 1A**). Injection of AAV2.1_CAGGS_Flex_ChR2_td-Tomato_WPRE_SV40 into GlyT2-Cre transgenic mice targeted GlyT2(+) cells (**Figure 1B**). Co-injection of AAV2.1-HSyn-Cre_WPRE_hGH with AAV2.1_CAGGS_Flex_ChR2_td-Tomato_WPRE_SV40 in GlyT2-eGFP mice enabled expression in all GoC subtypes (**Figure 1C**). In this second strategy, GlyT2(-) GoCs are identified by the absence of GFP expression (**Figure 1D**), which has been previously confirmed to coincide with an absence of GlyT2 transport current in this line (Rousseau et al 2008). GoCs were easily recognized by their large somas in the granular layer and characteristic apical dendrites in the molecular layer (**Figure 1B** and **D**).

**Figure 1.**
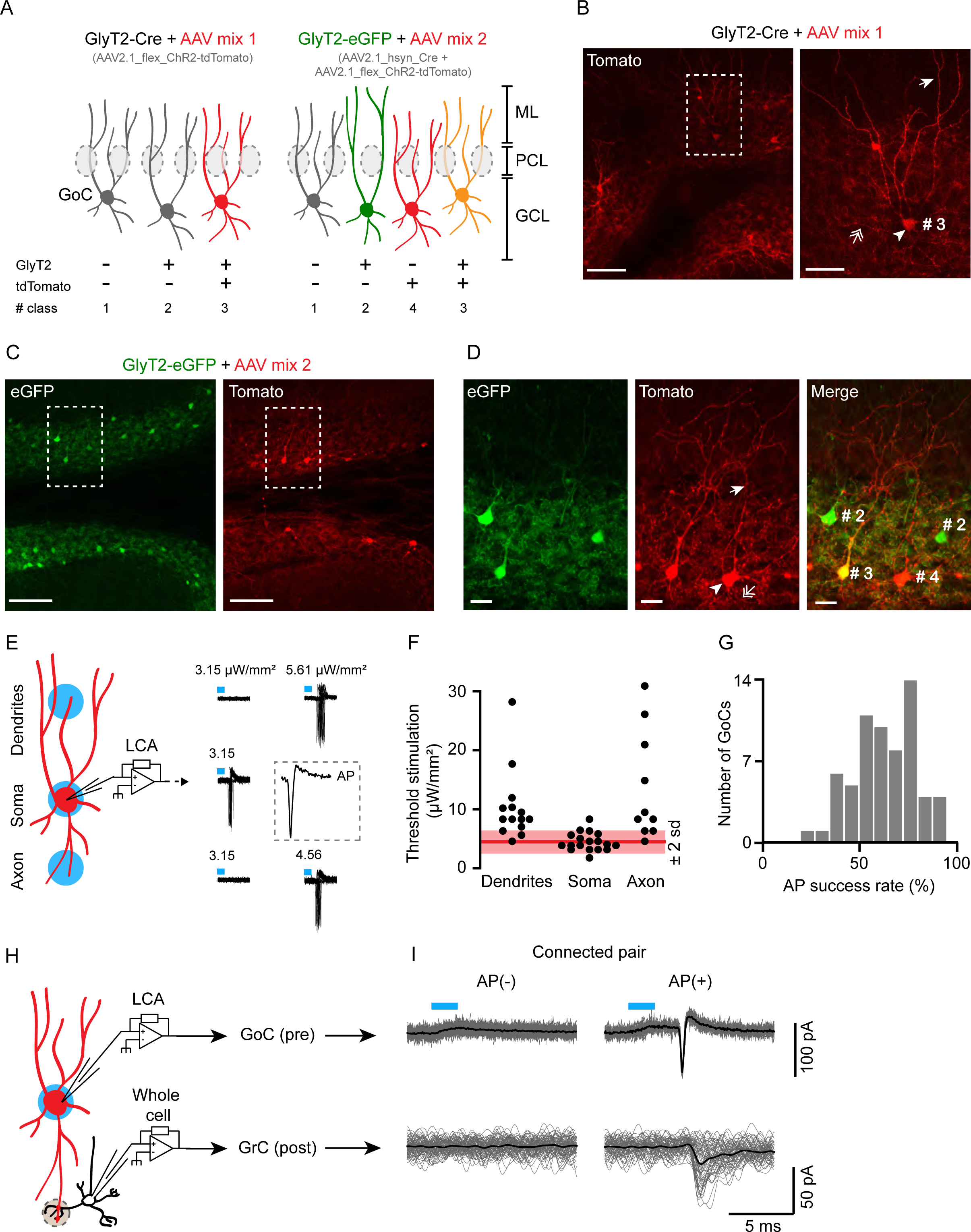
**A targeted optogenetic stimulation strategy for GoC-GrC paired recordings without presynaptic dialysis. A**, schematic of the viral strategies to express the channelrhodopsine (ChR2) fused with the tdTomato as reporter in GlyT2(+) and GlyT2(-) GoCs. GCL : granular layer, PCL : Purkinje cells layer, ML : molecular layer. **B**, slice of a representative GlyT2-Cre mouse injected with the mix 1 of AAV. Left panel, apex of the lobule IV showing infected, tdTomato(+) GoCs. Scale bar : 100 µm. Right panel, left panel magnification showing the characteristic GoCs morphology. Arrowhead : soma; arrow : apical dendrite; double arrow : axons. Scale bar : 20 µm. **C**, as B for a GlyT2-eGFP mouse injected with the mix 2 of AAV. Scale bar : 100 µm. **D**, magnification of C. (#2 and #3) : GlyT2(+) GoC; (#4) : GlyT2(-) GoC. Scale bar : 20 µm. **E**, left panel, calibration of the optogenetic stimulation power (in blue) to reach > 80 % of AP success (Threshold stimulation). Right panel, threshold stimulation power at the soma, dendrites, and axons of a representative GlyT2-Cre(+) GoC. Insert, magnification of an AP evoked at the soma. **F**, populational distribution of the threshold stimulation power (dendrites n = 13, soma n = 18, axons n = 10). In red, mean power at the soma ± 2 sd. **G**, Histogram of AP success rate evoked by somatic subthreshold stimulation in connected and unconnected GlyT2(+) and GlyT2(-) GoCs (n = 64). **H**, Schematic of the GoC-GrC optogenetic pairs recording of the GABAergic transmission in cerebellar glomeruli (grey dotted circle). **I**, representative example of a specifically connected GoC-GrC pair.

Optogenetic stimulations of GoCs using a restricted illumination field (20-30 µM in diameter) on the cell soma have previously been performed (Crowley et al 2009). However, the specificity of optogenetic stimulations needs to be validated, as dendrites and axons of other GoCs (ChR2+) could cross the stimulation light beam above or below the targeted soma, as illustrated by the extensive neuritic expression of td-Tomato(+) in the granular layer in our GoC expression paradigm (**Figure 1B** and **1C**). We therefore calibrated the optogenetic stimulation on slices of GlyT2-Cre animals expressing ChR2 in GlyT2(+) GoCs by viral transgenesis. We recorded GoCs in the cell-attached configuration and defined a stimulation threshold for each targeted cell as the lowest light intensity to evoke > 80 % of AP (**Figure 1E**). This stimulation threshold was always lower when the illumination spot was centered on the soma (4.47 ± 1.5 µW/mm², n = 18) than when placed remotely on the dendritic (10.3 ± 6 µW/mm², n = 13) or axonal fields (13.6 ± 8.8 µW/mm², n = 10) of the same GoC (soma vs. dendrites: p <0.001, soma vs. axon: p <0.001, Wilcoxon’s bilateral test). However, the power used for the least excitable cells at the soma could be supra-liminal for a small fraction of axons or dendrites of a neighboring highly excitable GoC (**Figure 1F**).

As there is a priori no guarantee that IPSCs evoked by efficient somatic stimulation originate exclusively from the target cell, we reduced and set the intensity of the optogenetic stimulation around the AP threshold (AP success rate : 62 ± 16 %, n = 64; **Figure 1G**). To systematically verify the specificity of this juxta-threshold optogenetic stimulation during paired recordings of GoCs and GrCs, we recorded light-evoked action potentials from the targeted GoC with a loose cell-attached patch pipette placed on its soma (**Figure 1H**), and we verified that AP failure in the GoC always resulted in IPSC failure in the postsynaptic GrC (**Figure 1I**).

### GlyT2(-) and GlyT2(+) GoC-GrC connections have different synaptic properties

To quantify the synaptic currents in GrC whole-cell recordings with maximum sensitivity, we calculated the 5-ms time integral of the current starting 1 ms after the GoC spike, and after subtracting the average current for the preceding 10 ms (**Figure 2Ai**). As expected, the integrated charge sampled randomly from the baseline before stimulation followed a very narrow distribution centered on zero (**Figure 2Aii-iii**, grey solid line, see technical details in materials and methods). In 26 of 64 pairs, the post-AP distribution of charge was not significantly different from the baseline distribution (**Figure 2Aii**), indicating unconnected pairs (Kolmogorov-Smirnov test p > 0.05), whereas the distribution was highly significantly right-shifted in the remaining 38 connected pairs (**Figure 2Aiii**, Kolmogorov-Smirnov test, p < 0.0001).

The mean amplitude of IPSCs, evoked by effective optogenetic stimulation that triggered an AP in the connected GoCs, has a large coefficient of variation (48.4 ± 37.8 pA, n = 38 pairs, CV = 0.78; **Figure 2B** and **C**), and shows distinct distributions for GlyT2(-) GoCs (70.3 ± 53.9 pA, n = 10) from GlyT2(+) GoCs (40.6 ± 26 pA, n = 28; Conover test of equal variance: p = 0.017; **Figure 2C**). Although the lower mean amplitude of GlyT2(+) GoCs IPSC is not statistically significant from GlyT2(-) GoCs IPSC (p = 0.28), the non-equality of variance is consistent with the hypothesis that the phenotypic diversity of GoCs alters the synaptic properties of the GABAergic synapses they form with GrCs.

**Figure 2.**
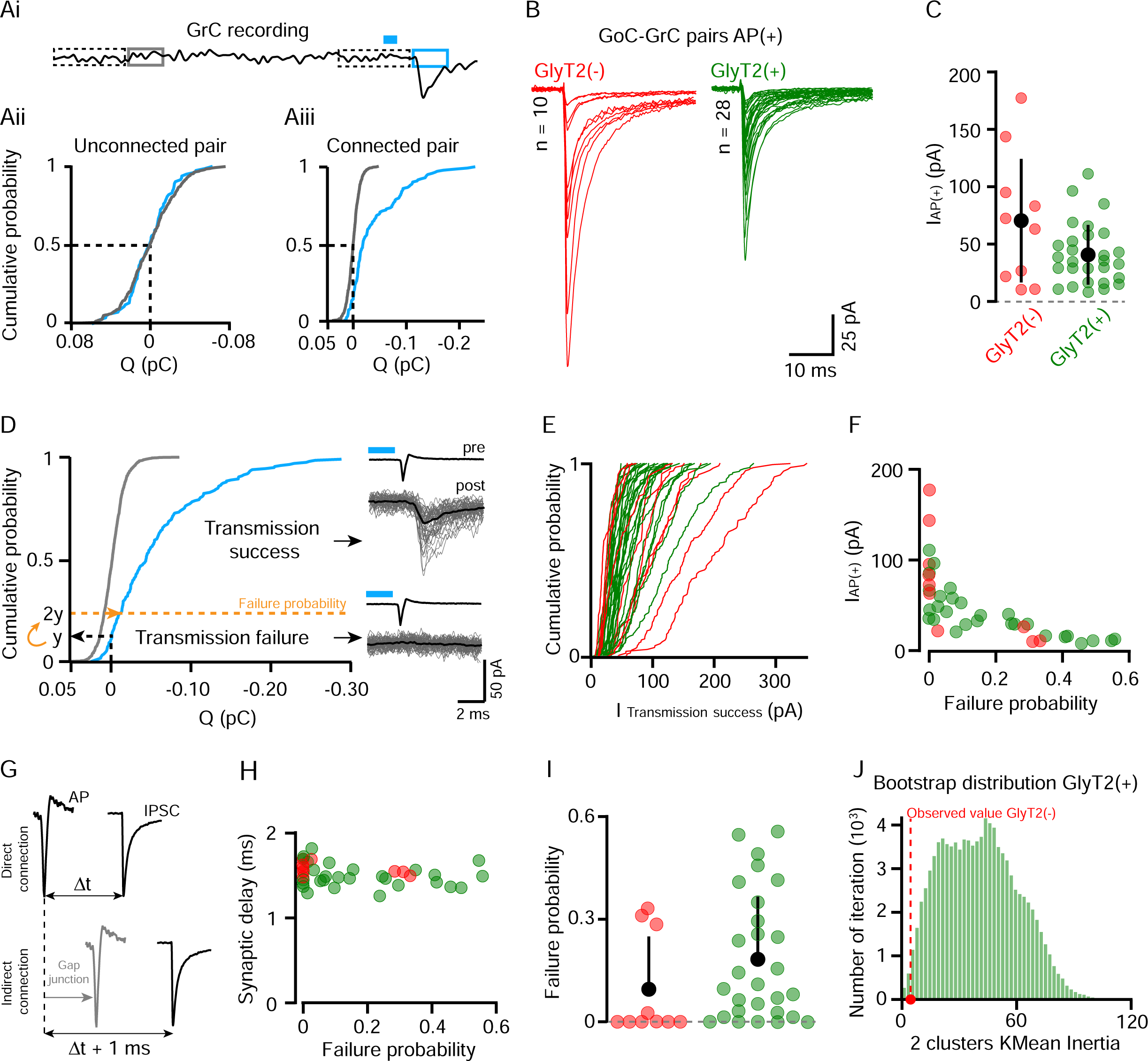
**GlyT2(-) and GlyT2(+) GoC-GrC connections have different synaptic properties. Ai**, illustration of the charge measurement in GrCs recordings before (grey) and after optogenetic stimulation evoking GoCs AP (blue). Dotted rectangles represent the local 10 ms normalization windows (see materials and method). **Aii**, Cumulative probability distribution of the baseline (grey) and post-AP (blue) charges from a representative unconnected pair. **Aiii**, Same as Aii for a representative connected pair. **B**, Average GrCs traces following optogenetically evoked AP in GyT2(-) (red, n = 10) and GlyT2(+) (green, n = 28) GoCs. These traces will be referred as AP(+) from here and IAP(+) is the peak amplitude of the averaged AP(+) traces. **C**, IAP(+) of GlyT2(-) and GlyT2(+) pairs shown in B. **D**, Same representation as Aii-Aiii, showing the transmission failure threshold of a representative connected pair (dotted yellow line). Right insert, resulting classification of the example traces in transmission success or failure with the corresponding average (black). **E**, Cumulative distribution probability of the transmission success peak amplitude for each GoC-GrC pairs. These curves show that GlyT2(-) pairs are more right shifted than the GlyT2(+) ones, but no significant differences emerge with this metric cleaned of transmission failure (GlyT2(-) : 80.78 ± 51.75 pA, n = 10; GlyT2(+) : 53.94 ± 23.75 pA, n = 28; p = 0.3). **F**, Transmission failure probability in function of the IAP(+) for all connected pairs (n = 38). **G**, Schematic illustrating the consequence of indirect GoC-GrC connection through Gap junction on the synaptic delay. **H**, Transmission failure probability in function of the synaptic delay (peak-to-peak time between AP and IAP(+); GlyT2(-) : 1.57 ± 0.072 ms, n = 10; GlyT2(+) : 1.49 ± 0.133 ms, n = 28; p = 0.04). **I**, Failure probability distribution showing a bimodal distribution for GlyT2(-) GoCs pairs and a continue one for GlyT2(+) ones. **J**, Histogram of the inertia for 2 clusters KMean analysis on bootstraped distribution of GlyT2(+) failure probability distribution (green). The red dot is the inertia of GlyT2(-) 2 clusters KMean analysis on the distribution shown in I. This figure shows how likely 8 points draw randomly with replace from the GlyT2(+) distribution give rise to a distribution as separated as the GlyT2(-) one.

### Different connectivity rules govern GlyT2(-) and GlyT2(+) GoCs synaptic contacts with GrCs

The transmission failure rate is an important quantal parameter of synaptic transmission, indicative of the number of active sites and the probability of vesicular release. The large difference between the synaptic and baseline charge distribution in connected pairs separated transmission successes from failures (**Figure 2D**). We set the postsynaptic failure probability for synaptic currents to twice the probability of measuring positive charge (outward current) (**Figure 2D**), knowing that the charge distribution of failures is symmetric around zero in the baseline (**Figure 2Aii-iii**). This limit defines a conservative estimate of the failure rate (0.16 ± 0.18, n = 38) since all traces below this threshold show no IPSC waveform. Eliminating transmission failure does not change the broad distribution of IPSC amplitude for each connected pair (p = 0.3), suggesting the contribution of multiple sources of variability (**Figure 2E**).

The occurrence of pairs with a high probability of transmission failure and a low amplitude IPSC (10 pairs out of 38 have a transmission failure probability between 0.3 and 0.55 in both GlyT2(-) and GlyT2(+) GoCs populations; **Figure 2F**) drew our attention because of a possible stimulation artifact that could confound our analysis. Indeed, it is well established that a fraction of the AP evoked in a GoC can propagate to neighboring GoCs through electrical gap junctions, emulating an indirect connection with low release probability (**Figure 2G**) (Dugue et al 2009, Vervaeke et al 2010). To rule out this interpretation, we measured the latency of evoked IPSCs with respect to the peak of the GoC AP, knowing that spike propagation *via* gap junctions introduces an additional delay of ∼1 ms (Dugue et al 2009, Vervaeke et al 2010). In our data, the mean synaptic delay had a clear monomodal distribution (1.51 ± 0.125 ms, n = 38), and did not correlate with the probability of transmission failure (r = -0.25, p = 0.11, n = 38, **Figure 2H**), confirming the pre-post synaptic specificity of our recorded pairs and the validity of our failure probability measurements. Failure rates were on average lower for GlyT2(-) pairs but not significantly different from GlyT2(+) ones (GlyT2(-) : 0.095 ± 0.14, n = 10; GlyT2(+) : 0.18 ± 0.18, n = 28; p = 0.061; **Figure 2I**).

However, K-means analysis shows that the failure rate of GlyT2(-) GoCs follows a bimodal distribution whereas this does not seem to be the case for GlyT2(+) GoCs (k = 2 clusters explain 99% and 80.5% of their dispersion, respectively). Bootstrap analysis performed by drawing 10 samples from the GlyT2(+) distribution and calculating their K-means inertia shows that the probability that the inertia of the two clusters from the GlyT2(-) distribution is explained by a different drawing of the GlyT2(+) distribution is 0.8 % (one-sided) (**Figure 2J**; see details in the methods). Together, these results (**Figure 2B-F**) show that GlyT2(-) GoCs follow different synaptic connectivity rules than GlyT2(+) GoCs.

### Different types of IPSCs unravel decreased and variable GABA transient at GlyT2(+) synapses

As the peak amplitude of the IPSCs is mostly affected by classical parameters defining the synaptic strength, such as the number of release sites and of postsynaptic receptors, it is not a sensitive readout of the potential impact of differential GABA contents in GoCs. We thus looked for other metrics (kinetics and charge) that may better detect changes in GABA content while reducing other potential sources of variability.

The decay of AP(+) current was well fitted by a bi-exponential function (unitary examples of each group are shown in **Figure 3A**), which yielded time constants characteristic of mature GoC-GC synapses (Tau1 : 1.97 ± 0.6 ms, Tau2 : 16.45 ± 6.26 ms, n = 36; **Figure 3B**) (Brickley et al 1996, Crowley et al 2009, Rossi & Hamann 1998). The two-time constants were not correlated across cells (Tau1 vs Tau2, r = 0.263, p = 0.12, n = 36; **Figure 3B**), and were undistinguishable between the two GoC types. As expected from the amplitude distributions, the mean AP(+) charge was 1.96-fold higher for GlyT2(-) than GlyT2(+) GoCs (0.4 ± 0.3 pC (n = 10) and 0.2 ± 0.13 pC (n = 26), respectively; p = 0.1; **Figure 3C**). Surprisingly, given the similar kinetic time constants in both GoC types, the QAP(+)/IAP(+) ratio, which establishes a mean Tau weighted, was 21 % higher in the GlyT2(-) group (GlyT2(-): 6.0 ± 0.82 ms, n = 10; GlyT2(+): 4.89 ± 1.5 ms, n = 26; p = 0.0154; **Figure 3D**) indicating a prolonged activation of GABAA receptors at GlyT2(-) synapses. This is explained by a larger contribution of the second component in the GlyT2(-) than GlyT2(+) population, as displayed in the mean traces of normalized IPSCs in the two populations(A2/A1+A2 : 0.281 vs 0.219, respectively, **Figure 3E**).

**Figure 3.**
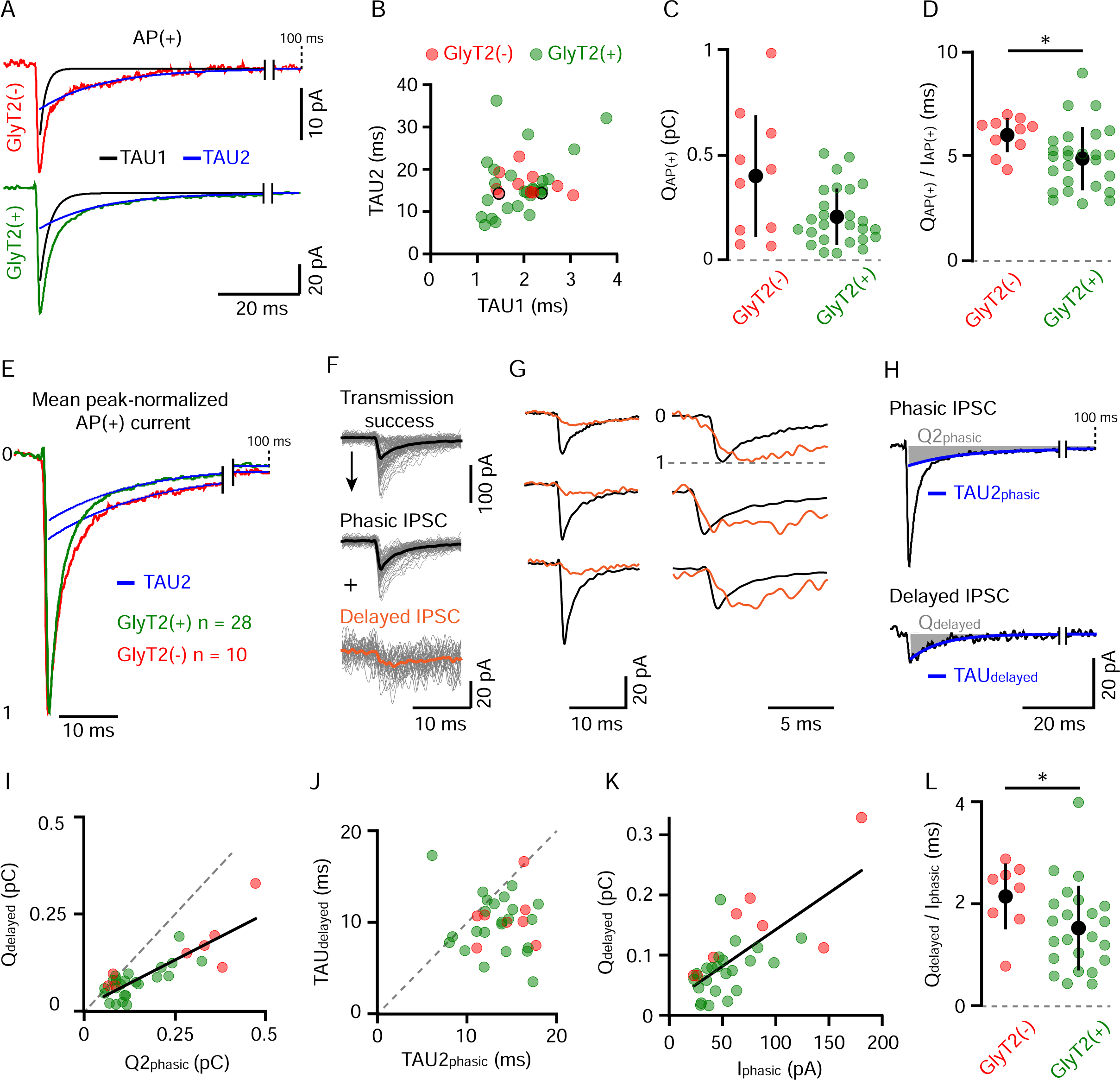
**Different types of IPSCs unravel decreased and variable GABA transient at GlyT2(+) synapses. A**, Example of GlyT2(-) and GlyT2(+) AP(+) averaged traces both adjusted by a bi-exponential function. The two decay components have a fast (black) and a slow (blue) time constant named TAU1 and TAU2 respectively. **B**, Correlation between the two fitted time constants in connected GoC-GrC pairs (n = 36, GlyT2(-) n = 10, GlyT2(+) n = 26). The two dots outlined in black are the individual examples in D. GlyT2(-) : TAU1 = 2.04 ± 0.51 ms, TAU2 = 16.56 ± 2.73 ms, n = 10; GlyT2(+): TAU1 = 1.94 ± 0.62 ms, TAU2 = 16.41 ± 7.17 ms, n = 26; TAU1 GlyT2(-) vs TAU1 GlyT2(+), p = 0.46, TAU2 GlyT2(-) vs TAU2 GlyT2(+), p = 0.58. **C**, Distribution of the synaptic charge (QAP(+)) calculated from bi-exponential fit (A1*TAU1+A2*TAU2) of mean AP(+) current. **D**, Ratio between QAP(+) and the peak amplitude of AP(+) current (IAP(+)), Figure 2B). **E**, Mean peak-normalized AP(+) current of the GlyT2(-) and GlyT2(+) pairs (n = 10 and n = 28 respectively). The superposed blue traces are the second decay component from the bi-exponential function (same fitting procedure as A). **F**, Transmission success from a representative pair decomposed in phasic (average trace in black) and delayed (average trace in orange) events (see details in materials and methods). **G**, Examples exhibiting kinetics difference of phasic and delayed events (left : average traces; right : peak-normalized traces). **H**, Average phasic and delayed IPSC of a pair adjusted with a bi- and a mono-exponential function reciprocally. In blue, the second decay component of the phasic IPSC (TAU2phasic) and the single decay component of the delayed IPSC (TAUdelayed). The areas shaded in grey represent the charge of the corresponding decay component in phasic (Q2phasic = A2*TAU2phasic) and delayed (Qdelayed = A*TAUdelayed) events. The peak amplitude of the phasic IPSC (Iphasic) did not differs significantly between GlyT2(+) and GlyT2(-) pairs. **I**, Correlation between Q2phasic and Qdelayed. **J**, Correlation between the decay time constant of delayed IPSCs (TAUdelayed) and the second one of phasic IPSCs (TAU2phasic). In I and J, the grey dotted lines are diagonals, and exhibit a lack of charges and time in delayed IPSCs compared to the phasic ones. **K**, Correlation between the Iphasic and Qdelayed (GlyT2(-) n = 8; GlyT2(+) n = 23). **L**, Qdelayed to Iphasic ratio for GlyT2(+) and GlyT2(-) pairs. These ratios are the slopes of each dot in K and depict that GlyT2(+) variability exceeds the one of GlyT2(-) pairs.

We thus examined further the origin of the second kinetic component of the IPSCs decay in Individual traces (see materials and methods). It revealed a subpopulation of low-amplitude, delayed IPSCs lacking the fast-rising fast-decaying component characteristic of phasic IPSCs (**Figure 3F-H**). These delayed IPSCs have similar prevalence in the two GoC populations (GlyT2(-) : 23.5 ± 25.5 % n = 10, GlyT2(+) : 26.2 ± 13.8 %, n = 28, p = 0.28) and their decaywas well fitted by a single exponential function (TAUdelayed : 10 ± 3 ms, n = 31). The charge of the delayed IPSCs was 2-fold higher in GlyT2(-) pairs than GlyT2(+) ones (Qdelayed GlyT2(-) : 0.147 ± 0.08 pC, n = 8; GlyT2(+) : 0.073 ± 0.04 pC, n = 23; p = 0.0176) and was highly correlated with the Q2phasic component (see **Figure 3H**) for each pair although smaller (slope = 0.48, r = 0.716, p < 0.0001, n = 31, **Figure 3I**). This reduction in Qdelayed was partly accounted by a faster decay (TAUdelayed vs TAU2phasic, p < 0.0001, n = 31, two-tailed Wilcoxon test, **Figure 3J**).

The fact that Qdelayed correlates with Iphasic (p < 0.0001, n = 31, **Figure 3K**), but is smaller than Q2phasic suggests that delayed IPSCs result from partial activation of the same clusters of synaptic receptors that mediate phasic IPSCs, but *via* a reduced and/or altered neurotransmitter transient (partly filled vesicle, distant release site, slow fusion…). To estimate the GABA transient evoking delayed IPSCs, as compared to phasic ones, we computed the Qdelayed/Iphasic ratio, which was found to be significantly higher for GlyT2(-) (2.1 ± 0.64 ms, n = 10) than for GlyT2(+) cells (1.5 ± 0.82 ms, n = 23; p = 0.032, **Figure 3L**). Altogether, the skew of the distribution of normalized QAP(+) and Qdelayed at GlyT2(+) synapses towards lower values, as compared to GlyT2(-) connections, indicate a reduced capacity to activate postsynaptic GABAAR with slow kinetics in a fraction of the GlyT2(+) GoCs, putatively as the result of a reduced synaptic GABA transient.

### Stronger activation of high-affinity GABAAR by GlyT2(-) GoCs

In addition to the classical fast IPSC, inhibitory transmission between GoCs and GrCs involves a slower component that is mediated by high-affinity extrasynaptic GABAARs activated by spillover and by accumulation of GABA in cerebellar glomeruli (Mapelli et al 2014, Rossi & Hamann 1998). In connected pairs, the time-integral of the average AP(+) traces (cumulated charge) revealed this slow component that prolongs the fast IPSC (**Figure 4A**), and which carries 2.5 ± 1.5 times more charges (0.62 ± 0.38 pC, n = 32, p < 0.0001, two-tailed Wilcoxon test).

**Figure 4.**
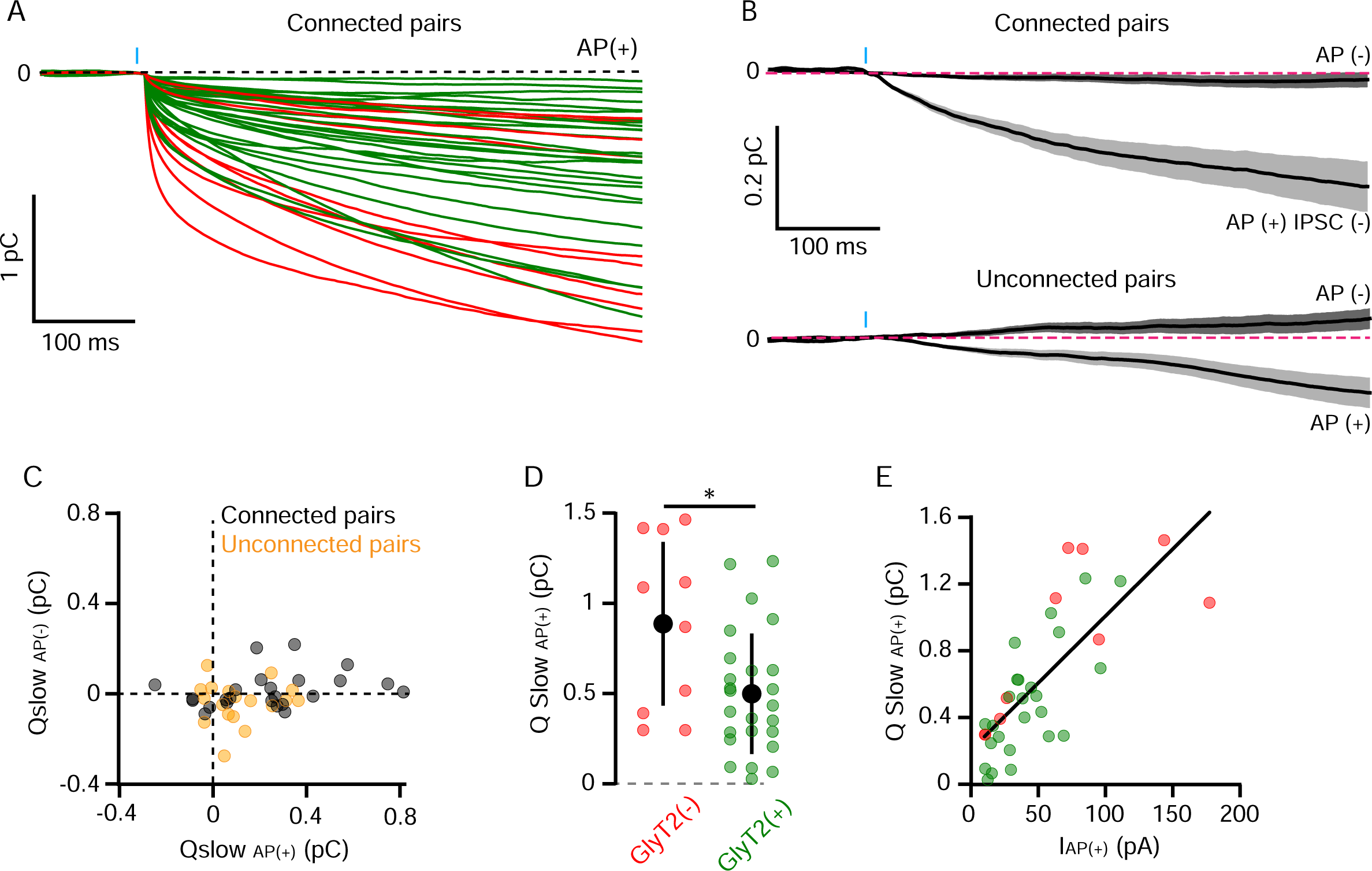
**A stronger activation of high-affinity GABAAR by GlyT2(-) GoCs. A**, Cumulative sum (integral) of the mean AP(+) current for each GoC-GrC pairs (red = GlyT2(-), n = 10; green = GlyT2(+), n = 27). **B**, Average integrals ± SEM for AP failure (AP(-)) or IPSC failure (AP(+) IPSC(-)) in connected (top, n = 24) and unconnected pairs (down, n = 18). **C**, Slow component charges (Qslow) in AP(-) and AP(+) IPSC(-) mean current from connected (black, n = 24) and unconnected pairs (yellow, n = 18). **D**, Qslow AP(+) in GlyT2(+) (n = 25) and GlyT2(-) (n = 10) pairs. **E**, Correlation between IAP(+) and Qslow AP(+).

In a subset of Goc-GrC pairs, the recording of 2 seconds post-AP allow us to fit the slow charge component (Qslow) with a mono-exponential function whose decay time constant (tauslow = 359 ± 111 ms (n = 6)) is in the range of previously recorded high-affinity extrasynaptic GABAAR components (Rossi & Hamann 1998). As expected, Qslow measured in traces without presynaptic AP did not differ from zero (0.009 ± 0.087 pC, n = 42), further confirming the specificity of our optogenetic pairs. In connected pairs, Qslow measured in the IPSC failure traces (AP(+) IPSC(-): 0.23 ± 0.25 pC) was smaller than in AP(+) IPSC(+) traces (0.6 ± 0.27 pC, n = 24, p < 0.0001, two-tailed Wilcoxon test), but significantly larger than in AP(-) : 0.0136 ± 0.077, n = 24, p = 0.0003, two-tailed Wilcoxon test; **Figure 4B** and **4C**). Finally, synaptic release of GABA from unconnected pairs also produced small but significant Qslow (AP(+) : 0.11 ± 0.13 pC; AP(-) : -0.039 ± 0.09 pC, n = 18, p = 0.004, two-tailed Wilcoxon test **Figure 4B** and **4C**). Overall, these data confirm that extrasynaptic high affinity GABAAR expressed by GrCs can be activated by spillover of GABA released from a single GoC after a single AP, even in the absence of a direct synaptic connection, but that most of their activation results from GABA release at synaptic contact sites. Given the transmitter pooling properties of cerebellar glomeruli (Brandalise et al 2012, Crowley et al 2009, DiGregorio et al 2002, Rossi & Hamann 1998), we reasoned that the slow component should be a good indicator of the vesicular release of GABA by stimulated GoC in the glomerulus. Remarkably, synaptic currents carry on average 1.77 times more slow charges when evoked by GlyT2(-) GoCs than by GlyT2(+) GoCs (GlyT2(-) : 0.88 ± 0.45 pC, n = 10, GlyT2(+) : 0.49 ± 0.33 pC, n = 25, p = 0.0298; **Figure 4D**). We found that the charge of the slow spillover component correlated with the amplitude of the phasic IPSC, IAP(+), across pairs (slope = 7.3 ms, r = 0.71, p < 0.0001, n = 31, **Figure 4E**) but the charge/amplitude ratio was not different between GlyT2(+) and GlyT2(-) GoCs (12.8 ± 7.4 and 17.5 ± 7.2 ms respectively; p = 0.082).

### IPSC modulations induced by manipulation of GABA and glycine supply to GoCs recapitulate GlyT2-linked IPSC variability

Our findings of a relationship between GlyT2 expression and a decrease in GABAergic transmission, particularly the slow kinetic component of the phasic IPSCs, could arise from the glycine accumulation activity of GlyT2, most likely through competition with GABA for vesicular loading, or from a spurious correlation with the expression of other molecular elements controlling synaptic wiring and synaptic transmission. We therefore sought to test directly in the slice whether altering the cytosolic glycine/GABA concentration ratio through increased GlyT2 activity could alter the strength of GABAergic inhibition, as previously demonstrated in cultured cells (Aubrey et al 2007, Rousseau et al 2008).

GABAergic IPSCs were evoked in GrCs by electrical stimulation of GoCs axons at 10 Hz (**Figure 5A**), a frequency similar to their average firing behavior in rodents *in vivo* (Holtzman et al 2006a, Simpson et al 2005, Vos et al 1999). Averaged IPSCs were computed every 100 stimulations (**Figure 5B**) and the charge to peak-amplitude ratio was monitored over time after normalization to baseline (**Figure 5C**). As suggested in Figure 3, this ratio may reflect the characteristics of the GABA transient activating the GABAAR of the recorded GrC. Bath application of glycine (100 µM) caused a gradual decrease in the charge to amplitude ratio, which did not appear to saturate even after 5 minutes of application (**Figure 5D**). Because transmission undergoes a slow rundown over time (see Material and Methods), controls without bath application of glycine were randomly performed. The charge to peak ratio was statistically smaller in the presence of glycine (t2 to t1 ratio in control: 0.91 ± 0.22, n = 10; glycine: 0.64 ± 0.18, n = 10, p = 0.017), revealing a 29 % glycine-induced decrease in charge to peak-amplitude ratio (**Figure 5E**). Furthermore, the depression of charge to peak-amplitude by glycine applied in the bath did not significantly reverse after 6 minutes of washout (t3 to t1 ratio in control: 0.83 ± 0.32, n = 8; in glycine: 0.46 ± 0.26, n = 10; p = 0.023; 44.6 % reduction) (**Figure 5E**).

**Figure 5.**
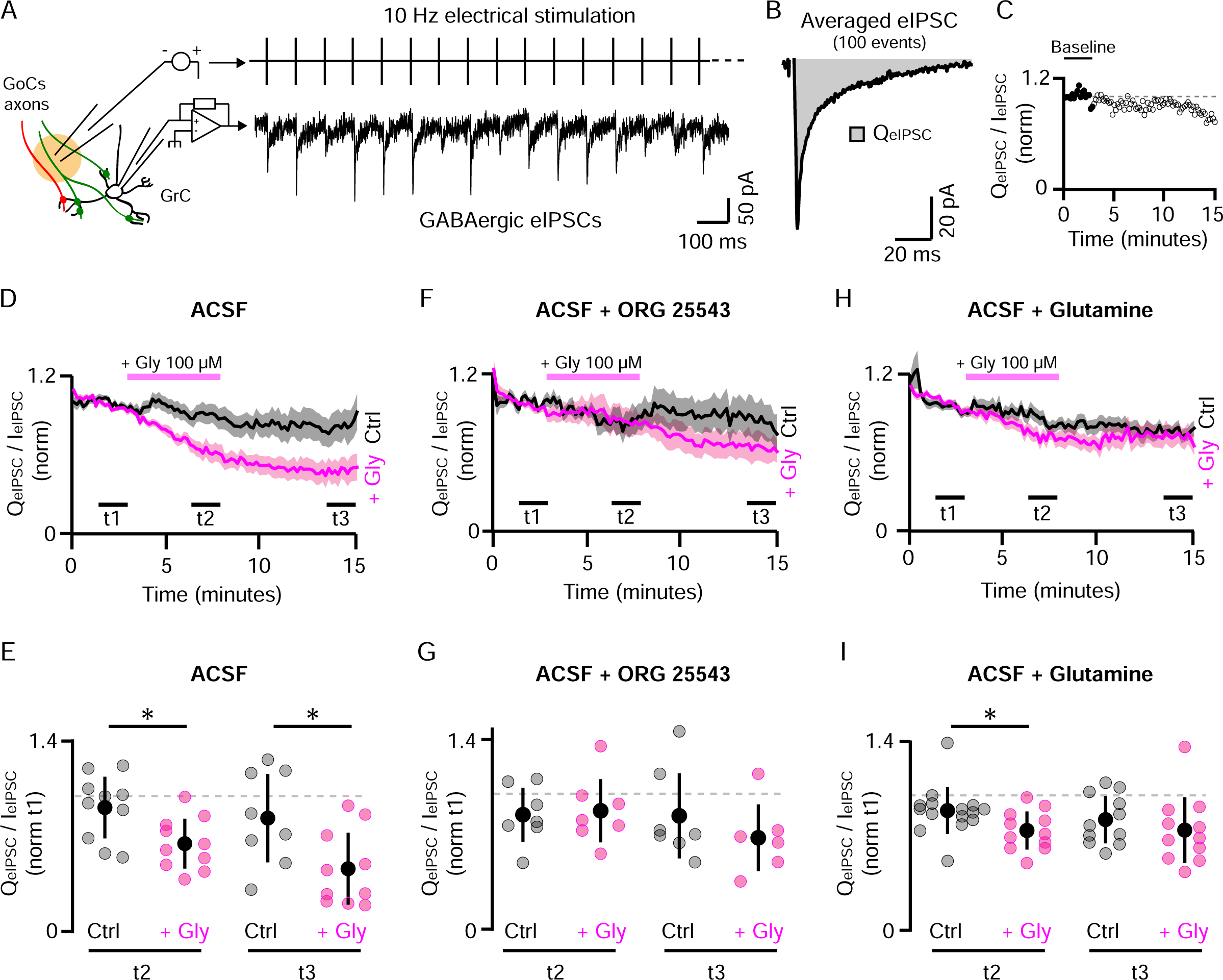
**manipulation of GABA and glycine supply to GoCs recapitulate GlyT2-linked IPSC variability**. **A**, left, schematic of the experiment. GABAergic IPSCs recorded from GrCs in whole cell configuration are evoked by the continue electrical stimulation (eIPSC) at 10 Hz of GlyT2(-) and GlyT2(+) GoCs axons without any distinction. Right, a GrC recording during 10 Hz stimulation of the GoCs axons. GABAergic eIPSCs have been pharmacologically isolated with the following cocktail of blockers : APV 50 µM, NBQX 2 µM, strychnine 0.5 µM, CGP55845 1 µM, ORG24598 1 µM. **B**, Average of 100 consecutive GrC eIPSCs following the electrical stimulation. The strength of the GABAergic transmission over the time is estimated by taking the charge (QeIPSC, grey area) to peak amplitude (IeIPSC) ratio of averaged eIPSCs every 10 seconds. **C**, Example of the QeIPSC to IeIPSC ratio over the time. Each dot was normalized by the mean signal in the baseline (3 minutes from the beginning, full dots). **D**, Populational evolution of the GABAergic transmission over the time in ACSF (ctrl, black, n = 8-10) or ASCF + glycine 100 µM for 5 minutes (+Gly, pink, n = 10). Traces represent the mean normalized charge to amplitude ratio ± SEM. **E**, Charge to amplitude ratio measured from 1000 consecutive eIPSCs averaged at the end of the glycine application (t2) and at the end of the recordings (washout, t3), normalized by the ratio at the end of the baseline (t1). These new quantifications were made from not normalized raw recording of each GrCs. **F-G**, same as D-E in presence of ORG25543 1 µM, the specific blocker of GlyT2 (F : ctrl n = 7, +Gly n = 6-7). **H-I**, Same as D-G in ACSF supplemented with 500 µM of Glutamine (Gln), the precursor of the GABA synthesis (H : ctrl n = 11-15, +Gly n = 11).

We monitored that the effect of glycine application was not due to a decrease in the efficiency of extracellular electrical stimulation caused by the activation of inhibitory glycine receptors (GlyR) on GoCs (Dumoulin et al 2001), despite the continuous presence of 0.5 µM strychnine in the bath to block GlyRs. The success rate of juxta-threshold electrical stimulation of GoCs axons, assessed by cell-attached somatic-recordings of AP, was not affected by glycine application in the presence of strychnine (control : 0.59 ± 0.06; glycine : 0.68 ± 0.1, p = 0.46, n = 6, two-tailed Wilcoxon test). We also controlled that the bath application of 100 µM of glycine had not direct impact on synaptic GABAAR by fitting the decay time constants of the IPSCs at t2 with or without glycine (control Tau1 : 3.48 ± 1.2 ms, Tau2 : 36.5 ± 7.7 ms, n = 32; glycine Tau1 : 3.19 ± 1.3 ms, Tau2 : 36.7 ± 8.3 ms, n = 29; Tau1 control vs glycine : p = 0.2, Tau2 control vs glycine : p = 0.79).

To confirm that the negative effect of glycine on GABAergic transmission of GrCs involves GlyT2, we added its specific blocker, ORG25543 (1 µM), during pre-incubation of slices and continuously during the experiment. Under this condition, glycine application did not cause a reduction in the charge to peak-amplitude ratio, compared to the control without glycine application (**Figure 5F**) (t2 to t1 ratio in control: 0.84 ± 0.19, n = 7; in glycine: 0.87 ± 0.23, n = 7, p = 0.949; t3 to t1 ratio in control: 0.83 ± 0.31, n = 7; in glycine: 0.67 ± 0.24, n = 6; p = 0.474; **Figure 5G**).

To validate that the effect of glycine was due to presynaptic competition with GABA, we sought to increase its metabolic supply by providing extracellular glutamine, which is transported by neurons and used as a direct precursor of glutamate for GABA synthesis (Ishibashi et al 2013, Wang et al 2013). In ACSF supplemented with 500 µM glutamine (**Figure 5H**), extracellular application of glycine caused significant decrease in transmission but with 1.7 fold smaller magnitude than in not supplemented ACSF (t2 to t1 ratio in control: 0.88 ± 0.17, n = 15; glycine: 0.73 ± 0.14, n = 11, p = 0.025, 16.7 % of decrease, **Figure 5I**). Furthermore, the effect reversed to non-significant levels after 6 minutes of glycine washout (t3 to t1 ratio in control : 0.82 ± 0.17, n = 11, glycine: 0.74 ± 0.24, n = 11, p = 0.264), as observed for vesicular load in studies using whole-cell manipulations of cytoplasmic neurotransmitter content (Apostolides & Trussell 2013).

In conclusion, presynaptic glycine accumulation by GlyT2 may interfere with presynaptic GABA supply for GABAergic transmission at the GoC-GrC synapse in the adult cerebellum. The negative effect of glycine on the eIPSC charge/amplitude ratio is consistent with GABA/glycine competition for vesicular filling and provides a reasonable explanation for the IPSC charge/amplitude ratio variability in the GlyT2(+) population (**Figure 3D** and **3L**).

## Discussion

In this paper we show that the neurochemical diversity of GoCs correlates with the properties of the IPSCs they evoke in GrCs. We find that pure GABAergic GlyT2(-) GoCs exhibit a bimodal distribution of synaptic strength across GrCs with some very large connections, while GlyT2(+) GoCs exhibit a narrower unimodal distribution. In contrast, while GlyT2(-) GoCs IPSCs display a narrow distribution of their charge to peak ratio, GlyT2(+) GoCs IPSCs show a widespread distribution which is due to a variable amplitude of the second kinetic component of the IPSCs. This charge to peak modulation is reproduced by artificial manipulation of the GABA and glycine supply to GoCs, hinting for a presynaptic origin and consistent with GlyT2-mediated glycine accumulation playing an antagonist role on GABAergic transmission at the GoC to GrC synapses. Here we discuss the mechanisms and functional consequences of synaptic diversity in GoCs and argue for the recognition of GlyT2(-) GoCs as a population functionally separate from the GlyT2(+) GoCs.

### Regulation of the charge content of IPSCs at GlyT2(+) GoCs synapses by a presynaptic mechanism

The level of vesicular loading by GABA and glycine has been shown to control the level of postsynaptic receptors activation during mixed inhibitory transmission probed in cultured neurons or reconstituted systems (Aubrey et al 2007, Ishibashi et al 2013). In these systems, high levels of glycine accumulation are required to compete with GABA for vesicular loading, probably to compensate for the higher affinity of the vesicular transporter VIAAT for GABA (Aubrey et al 2007, Farsi et al 2016, McIntire et al 1997). This competition for VIAAT uptake has also been quantitatively assessed by directly manipulating cytoplasmic neurotransmitter content in acute slices and measuring mixed IPSCs amplitudes (Apostolides & Trussell 2013). Here, we propose that the charge-to-amplitude ratio of purely GABAAR-mediated GrCs IPSCs is a marker of the vesicular GABA released by the presynaptic element. Indeed, we show, by increasing extracellular glycine or extracellular glutamine, that the charge-to-amplitude ratio is acutely modulated by the competition between GABA and glycine availability for vesicular filling in GoCs.

IPSC kinetic modulation by cotransmission has previously been observed at MNTB synapses for instance, where GABA acts as a low affinity agonist on glycine receptors and accelerates the decay of glycinergic IPSCS (Lu et al 2008). Our results argue for a different mode of action, where a silent neurotransmitter (glycine) acts through vesicular filling competition with a postsynaptically active one (GABA).

### GlyT2(-) GoCs as a separate functional subtype mediating granular layer spatiotemporal patterning

Although we cannot rule out a marginal role of increased vesicular GABA content in the higher amplitude of GlyT2(-) pairs, the main explanation most likely lies in the specific synaptic connectivity rules between GlyT2(-) GoCs and GrCs, where GlyT2(-) GoCs make either small connections with high failure rate or very large connections with negligible failure. The specific patterns of strong synapses formed by each GlyT2(-) GoCs on a subsets of GrCs would be ideally suited to create spatial and temporal inhibitory patterning in the granular layer (D’Angelo et al 2013, Duguid et al 2015, Mitchell &Silver 2003). We therefore propose here that GlyT2(-) GoCs constitute a functional subtype, distinct from the rest of the GoC population. This is consistent with previous findings indicating that GlyT2(-) GoCs are specifically contacted by mixed inhibitory neurons from the cerebellar nuclei, which can control their firing (Ankri et al 2015). In this view, spatiotemporal patterning of the GrC population could therefore arise from disinhibition of granule cells by mixed inhibitory neurons from the cerebellar nuclei.

### Co-regulation of molecular and neurotransmitter phenotypes

The molecular diversity of GoCs extends beyond their neurotransmitter content, with the expression of various markers that appears to be co-regulated with the neurotransmitter phenotype. For instance, mGluR2 is expressed exclusively by GlyT2(+) GoCs (Simat et al 2007), while mGluR1/5 is expressed by a small population of GoCs that is similar in abundance to GlyT2(-) cells and does not overlap with mGluR2 expression (Negyessy et al 1997, Neki et al 1996). Serotonin receptors are also expressed by a subpopulation of GoCs and are able to increase the level of activity in the GoC network (Fleming & Hull 2019). Current single cell transcriptomic data have revealed a high degree of molecular diversity within cell types (Scala et al 2020), including cerebellar GoCs (Kozareva et al 2021). Single-cell transcriptomic data could be used to identify co-regulated molecular modules and highlight physiological properties underlying internal cell class diversity (Nusbaum et al 2017), such as the putative mGluR1/5-mGluR2-GlyT2-GAD module in GoCs. However, the presents results argue for the need of detailed physiological studies to test whether molecular diversity can be interpreted as a continuum of cell properties within a class or as the substrate to define cell subclasses (Petilla Interneuron Nomenclature et al 2008, Saunders et al 2018, Tasic et al 2016, Tremblay et al 2016, Yuste et al 2020, Zeisel et al 2015, Zeng & Sanes 2017).

### A push-pull hypothesis of gain control by mixed GABA/glycine inhibitory networks

The metabotropic mGluR2 operates a major inhibitory control of GoCs through massive activation of GIRK potassium channels (Watanabe & Nakanishi 2003). GoCs mGluR2 are recruited in a graded manner by inputs from mossy fiber (MF) and parallel fiber (Nietz et al 2017, Watanabe & Nakanishi 2003). The mGluR2 receptors can also be activated by neighboring climbing fibers through glutamate spillover, with GoCs displaying varying degrees of inhibition (Nietz et al 2017). Therefore, *in vivo*, sensory stimuli can cause long pauses in spontaneous firing of many GoCs (Holtzman et al 2006b, Holtzman et al 2011), in part due to activation of mGluR2 (Holtzman et al 2011). Overall, mGluR2 receptors act as a global sensor of excitatory cerebellar cortex activity in GlyT2(+) GoCs, which might be opposed by mGluR1/5 in GlyT2(-) GoCs expressing these receptors. It is therefore reasonable to propose that GoCs expressing high levels of mGluR2 could be arrested, rather than recruited, by an increased level of excitatory activity.

Based on our results on the variable synaptic strength of GoC connections, a preferential recruitment of glycine-rich cells at low levels of cerebellar activity and of GABA-rich cells at higher levels of activity would supra-linearly tune the level of inhibitory control over GrCs as a function of the overall level of MF and GrC activity, as GABA-rich cells produce large spillover and buildup components which can add up (Crowley et al 2009, Rossi & Hamann 1998). This organization is well suited to control the input-output relationship of the granular layer over a wide range of MF input activity (Mitchell & Silver 2003).

Glycine may also play an opposing role in this gain control scheme, as a co-agonist of NMDA receptors (Johnson & Ascher 1987). Potentiation of NMDA receptors by synaptically-released glycine has been demonstrated in the spinal cord (Ahmadi et al 2003). Given the low levels of D-serine in the adult cerebellar cortex (Koga et al 2017, Wang & Zhu 2003, Wolosker et al 1999) and the tight control of glycine extracellular levels by GlyT1 and GlyT2 transporters (Supplisson & Bergman 1997), both of which are present around and inside the cerebellar glomeruli (Zafra et al 1995a, Zafra et al 1995b), glycine released at GoCs synapses is a likely source of co-agonist for GrCs NMDA receptors. GrCs specifically express NR2C-containing NMDA receptors (Akazawa et al 1994, Cathala et al 2000, Farrant et al 1994, Monyer et al 1994) that are involved in the integration of MF input over long time scales (Baade et al 2016, Powell et al 2015, Schwartz et al 2012). This integration is greatest at low MF firing rates but can saturate at high MF firing rates. A decrease in glycine-rich GoC activity during high MF activity could decrease extracellular glycine and reduce NMDA excitation of GrCs, thereby increasing the integration bandwidth of the granular layer. This NMDA/GABA push-pull action, combined with the neurochemical diversity of inhibitory populations, could be a major mechanism in the lower brain to adapt, where needed, local circuit gain to the level of input activity.

## Materials and methods

### Animals

The experiments were performed on GlyT2-eGFP (Zeilhofer et al 2005) and GlyT2-Cre transgenic mice (kind gift of HU Zeilhofer, University of Zurich) of both sexes. For the optogenetic paired recordings, GlyT2-Cre and GlyT2-eGFP mice of 6-8 weeks have been used. For the pharmacological experiments, GlyT2-eGFP mice of 5-8 weeks have been used. Mice are derived and maintained on a C57BL6/j genetic background in our animal facility. All animal manipulations were made in accordance with guidelines of the Centre national de la recherche scientifique and Use Committee. This study was carried out under the ethical project number 02235.02.

### Stereotaxic injection

For GoC-GrC optogenetic paired recordings, cerebellar lobule IV/V of 4-5 weeks old GlyT2-Cre and GlyT2-eGFP mice were respectively injected with the adeno-associated viruses Mix 1 or 2, to infect GlyT2(+) GoCs or GlyT2(+) and GlyT2(-) GoCs. Mix 1 : AAV2.1_CAGGS_Flex_ChR2_td-Tomato_WPRE_SV40 (titration of 1.3-5.9*10^12, Upenn Vector Core, AV-1-18917), Mix 2 : AAV2.1_HSyn-Cre_WPRE_hGH (titration of 1.9-3.15*10^11, Upenn Vector Core, AV-1-PV2676) + AAV2.1_CAGGS_Flex_ChR2_td-Tomato_WPRE_SV40 (titration of 5.9*10^11, Upenn Vector Core, AV-1-1-18917). Mice are injected with Buprenorphine at 0.1 mg / Kg, 20-30 minutes before the start of the procedure. The animals are then induced with an Isoflurane / O2 mixture for 4 minutes at 3 % and kept under anesthesia for the duration of the procedure around 1.5-2 %. The correct placement of the head in the stereotaxic frame is confirmed by measuring a bregma-lambda Z deviation between 0.01 and -0.01 mm. A wide trepanation is then performed at -5.4 mm from the bregma, which reveals the bone thickening separating the colliculus from the cerebellum and the difference in contrast marking the transition between lobules III and IV. These internal parameters allow, if necessary, to adjust the anterior-posterior coordinates of the injection site which are likely to vary from one mouse to another at this age. Injections were performed with quartz (Sutter instrument, Novato, USA) or borosilicate (Harvard apparatus, Holliston, USA) capillary (length : 75 mm, O.D : 1 mm, I.D : 0.5 mm,) filled with Mix 1 or the Mix 2. GlyT2-Cre mice received one injection (anterior-posterior : -6 mm, medio-lateral : 0 mm, dorso-ventral : -0.300 mm) and GlyT2-eGFP received two medio-lateral injection to optimize Mix 2 virus expression (anterior-posterior : -6 mm, medio-lateral : 0 mm ± 0.250 mm, dorso-ventral : - 0.300 mm). Once in the tissue, the capillary is held for 2-3 minutes, then, 500 nl of virus per injection site are inoculated at constant speed (100 nl / min) and constant pressure using a Hamilton mounted on an injector (Hamilton, Reno, USA). At the end of the injection, the capillary is maintained for 10 minutes to let the liquid diffuse into the nervous parenchyma and then removed carefully. The transgenes were let to be expressed for 2 weeks before the experiment.

### Cerebellar slices preparation

All electrophysiological experiments were performed on 300 µm tick parasagittal slices of adult mice cerebellum following the same preparation procedure. After deep anesthesia induced with isoflurane (mix with 4 % medical oxygen in an induction box), mice were decapitated and the cerebellum was rapidly removed and dissected in a 4°C bicarbonate buffered solution (BBS, referred as ACSF in the results) containing the following (in mM) : 125 NaCl, 3.5 KCl, 1.25 NaH2PO4, 26 NaHCO3, 25 D-glucose, 1.6 CaCl2 and 1.5 MgCl2 (oxygenation 95% O2, 5% CO2). Slices are then cut using a vibrating blade microtome (Campden instrument (Loughborough, UK) or Leica VT 1000-S (Nanterre, France)) in a potassium-rich solution (GCS) at 4°C containing the following (in mM): 130 K-gluconate, 15 KCl, 0.05 EGTA, 20 HEPES and 25 D-glucose, the pH being adjusted to 7.4 by NaOH. The slices are then transiently immersed in a modified mannitol-based recovery solution (MRS) containing (in mM): 225 D-mannitol, 2.5 KCl, 1.25 NaH2PO4, 25 NaHCO3, 25 D-glucose, 0.8 CaCl2 and 8 MgCl2 (34°C, oxygenation 95% O2, 5% CO2) to help the gradual rebalancing of the ions towards normal external concentrations. 2-amino-5-phosphonovaleric acid (D-APV, Hellobio, Dunshaughlin, Republic of Ireland) at 50 µM was added to GCS and MRS to prevent glutamate excitotoxicity. Finally, the slices were transferred to oxygenated BBS (34°C, oxygenation 95% O2, 5% CO2) in which they were stored for a maximum of 6 hours. All components of BBS, GCS and MCS were purchased from Sigma Aldrich (Saint-Louis, USA) and the CaCl2 and MgCl2 from Fluka (distributed by VWR, Radnor, USA). BBS (ACSF), GCS and MCS are made in Volvic water (distributed by Danone, Paris, France) as previously detailed (Ankri et al 2015).

### Electrophysiology

#### Recordings

Prior to their electrophysiological recording, slices are transferred to a recording chamber mounted on an upright microscope (Olympus, Tokyo, Japan) and perfused (4 ml / min) with oxygenated BBS (95% O2, 5% CO2) at a temperature of 32-34°C under the objective. The slices are visualized thanks to an infrared-light source, a 20 X immersion objective (XLUM Plan FI, Olympus) and a camera (Cool Snap HQ, Photometrics, Tucson, USA). The recording and stimulation pipettes were stretched from borosilicate glass capillaries (length: 75 mm, outer diameter: 1.5 mm, wall thickness: 0.225 mm, Hilgenberg, Malfsfeld, Germany) with a vertical stretcher (David Kopf Instruments, Tujunga, USA). GoCs are identified in slices by the expression of GFP in GlyT2-eGFP and/or Tomato in injected mice. GrCs are easily recognized by their size, fast mono-exponential capacitive current and a capacitance < 4 pF (Silver et al 1992). The effect of optogenetic stimulations on targeted GoCs are recorded in loose-cell-attached and voltage clamp mode (holding at 0 mV) with 3-4 MΩ electrodes filled with an intrapipette solution containing (in mM): 140 NaCl, 2.34 KCl, 1.25 NaH2PO4, 10 HEPES, 1.3 CaCl2, 1.1 MgCl2, pH adjusted to 7.4 with 1M NaOH. For all experiments, GrCs are recorded in whole cell configuration and voltage clamp mode (holding at -70 mV) with 7-8 MΩ electrodes filled with an intracellular solution containing (in mM): 110 CsCl, 20 TEA-Cl, 10 HEPES, 6 NaCl, 10 EGTA, 0.2 CaCl2, 4 ATP-Mg, 0.4 GTP-Na, pH adjusted to 7.4 with CsOH at 1M. The data were acquired with an EPC10 amplifier (HEKA, Lambrecht, Germany), sampled at 20 KHz and filtered at 8 or 3 KHz. Chemicals for the internal solutions are from Sigma-Aldrich.

#### Optogenetic stimulation of GoCs

The source of light for optogenetic stimulations was a 470 nm LED (M470F3, THORLABS, Newton, USA) relayed to the sample by a collimation lens system and a lateral port of the Olympus equipped with the proper dichroic mirror. This optical setup allows to create a near-collimated spot of 20-30 µm in diameter on the sample. The optogenetic stimulation was delivered in 2 ms flashes which intensity was controlled linearly by the amplitude of the analog voltage step generated by the EPC10. The threshold for optogenetic stimulation intensity to GoCs was explored manually using a LEDD1B controller (THORLABS). The frequency of the optogenetic stimulation for GoC-GrC pair recordings was 0.37 Hz and 0.21 Hz when 2 additional seconds post-optogenetic stimulation were recorded (6/38 pairs).

#### Electrical stimulation of the GoCs axons

A current generator (isostim™ A320, WPI, Sarasota, USA) driven by the EPC10 amplifier has been used to deliver minimal electrical stimulations of 0.3 ms to the GoCs axons in GlyT2-eGFP slices. The stimulation electrode (7-8 MΩ) was filled with the same solution as for the loose-cell-attached recordings. The stimulation electrodes were systematically placed 50-100 µm away from the recorded GrC to avoid direct stimulation. The stimulation frequency was set at 10 Hz in continue for vesicular content manipulation experiments.

### Pharmacology

For the recording of the GoC-GrC optogenetic pairs, D-APV 50 µM, NBQX 2µM (Hellobio) and strychnine 0.5 µM (Sigma Aldrich) was added to the ACSF to avoid uncontrolled activation and inhibition of GoCs and isolate pure GABAergic IPSCs in our high chloride recording condition. The GABAergic IPSCs evoked in GrCs by the repetitive electrical stimulation of the GoCs axons have also been isolated by adding D-APV 50 µM, NBQX 2 µM, strychnine 0.5 µM and CGP55845 1 µM (Abcam, Cambridge, UK) to avoid GABABR activation in this condition (Mapelli et al 2009). In some experiments, the ACSF was supplemented with 500 µM of glutamine (Sigma Aldrich) to increases *de novo* synthesis of GABA (Wang et al 2013). Continuous repetitive 10 Hz stimulations of GoCs in adult slices led to rundown of the GABAergic transmission after 10 minutes of recording in classical slices preparation (39 ± 24 % of loss; n = 12). To stabilize the synaptic transmission during 10 Hz train stimulation, we improved the supply of essential substrates for general neuronal metabolism by incubating slices for at least one hour in BrainPhys (StemCell, Vancouver, Canada), a specially adapted supplemented culture medium (Bardy et al 2015). In these conditions, the rundown was reduced to 20 ± 27 % (no BrainPhys incubation (n = 12) vs BrainPhys incubation (n = 22) : p = 0.015). Due to the remaining rundown of the transmission over the time and its cell-to-cell variability, we randomly apply glycine (100 µM, 5 minutes, Sigma Aldrich) after 3 minutes of baseline recording. To avoid the buffering of the applied glycine by glial cells and that they became an uncontrolled source of glycine during washout, the glial transporter of glycine (GlyT1b) was blocked with of 1 µM of ORG24598 (Tocris, Bristol, UK) during the recovery and the experiment (Brown et al 2001). In other experiment, the neuronal transporter of glycine (GlyT2) has been blocked by adding 1 µM of ORG25543 (Tocris) (Caulfield et al 2001) during the recovery and the experiment. All drugs were applied in the recording chamber *via* the infusion system at the same rate and temperature than ASCF (4 ml/minute, 32-34°C).

### Image acquisition

The images of the infected and recorded slices were all acquired with an inverted confocal microscope equipped with a white laser (SP8, Leica).

### Analysis

#### Events detection and classification

The recordings were analyzed with algorithms developed on Python (Python Software Foundation, version 2.7). The method to differentiate connected from not connected pairs and transmission successes from failures are detailed in the text. Transmission successes are further segregated in phasic and delayed IPSCs based on the lack of the fast-rising and fast-decaying component in the latter. The difference in the rising phase kinetic has been measured with a sliding difference between the mean of a 10 ms time window and the mean of a 0.5 ms one, both separated by 0.3 ms corresponding to the rising time of a classical fast IPSC. The time to the maximal difference for a pair was set as the fast component rising time. Then, a 0.5 ms jitter of GoC-GrC transmission delay (Dugue et al 2005) was added around the fast component rising time previously measured to create, for each pair, the time window in which all fast-rising IPSCs should fall. An IPSC success was classed as phasic when the maximal difference in that window was superior to the baseline + 2 sd, otherwise it was classed as delayed IPSCs. The kinetics analysis of the IPSCs decay time were performed on Clampfit (Molecular Devices, San Jose, USA) from averaged traces.

### Statistics

The statistics were made with the scipy library available on Python (Virtanen et al 2020). The data are presented as mean ± sd unless otherwise stated in the text. All population difference significance was assessed by a bilateral non-parametric Mann-Whitney ranked test due to the small size of our groups and their no monotone distribution. Paired comparisons were calculated by a non-parametric Wilcoxon bilateral test when appropriated. All correlations have been calculated with the Spearman method and the corresponding two-sided p value were calculated with a t statistic. When correlations are significant, they are represented with the linear regression on the figure and the slope is reported in the text. The significance of the variance difference between GlyT2(+) and GlyT2(-) pairs have been tested by running a Conover test of equal variance (Wolfram function on Mathematica 12). An effect is considered significant if the p-value is less than 0.05. A p value is considered as strong when below 0.0001 and is reported as < 0.0001 in the text for clarity.

### Sample size estimation

In accordance with previous studies using cerebellar and brain stem slices (Ankri et al 2015, Apostolides & Trussell 2013, Bright et al 2011, Crowley et al 2009, DiGregorio et al 2007, Fleming & Hull 2019, Hirono et al 2012, Schonewille et al 2021, Stell et al 2003, Szoboszlay et al 2016) and the central limit theorem from the high number theory, we tried to reach a number of independent biological replicates (all “n” in this study) around 10 to 30 in each experiments.

### Attrition

Instable recordings from GrCs and spontaneous activity in GoCs were the only exclusion parameters used in this study. For the analysis of the optogenetic pairs, group size change along the study because the amount of recording in different class of event varies between pairs. Seven pairs (2 GlyT2(-) and 5 GlyT2(+)) did not have enough delayed IPSCs to be properly fitted with a mono-exponential function. Two GlyT2(+) pairs had a too small signal-to-noise ratio to be properly fitted with the two exponential function and could not be used for the quantification of QAP(+) and QslowAP(+). Another GlyT2(+) pair shows a marked artifactual rupture in its AP(+) current integral after 100 ms and were thus removed from QslowAP(+) analysis.

## Acknowledgments

We thank Boris Barbour, Johnathan Bradley, Mariano Casado and Vincent Villette for discussion on the manuscript and suggestion for the data analysis. DD was financed by a MESRI PhD fellowship, Labex memolife and ANR grant Glubrain3A. This study was supported by the grants équipe FRM DEQ20140329498 and ANR-17-CE16-014-03 GluBrain3A, by CNRS and by INSERM. We gratefully acknowledge the IBENS imaging facility (IMACHEM-IBiSA), member of the French National Research Infrastructure France-BioImaging (ANR-10-INBS-04), which received support from the “Fédération pour la Recherche sur le Cerveau - Rotary International France” (2011) and from the program « Investissements d’Avenir » ANR-10-LABX-54 MEMOLIFE. We also thank the teams of the animal facility and acute transgenesis facility (PFL2) of IBENS.

## Competing interests

SD is a stakeholder of the SME Karthala System and an inventor of IP licensed to this company.

## Data availability and code

Data and code for the electrophysiological analysis are available from the corresponding authors upon reasonable request.

## Notes

### Competing Interest Statement

The authors have declared no competing interest.

## References

1. Ahmadi S, Muth-Selbach U, Lauterbach A, Lipfert P, Neuhuber WL, Zeilhofer HU. 2003. Facilitation of spinal NMDA receptor currents by spillover of synaptically released glycine. Science 300: 2094–7

2. Akazawa C, Shigemoto R, Bessho Y, Nakanishi S, Mizuno N. 1994. Differential expression of five N-methyl-D-aspartate receptor subunit mRNAs in the cerebellum of developing and adult rats. The Journal of comparative neurology 347: 150–60

3. Ankri L, Husson Z, Pietrajtis K, Proville R, Lena C, et al. 2015. A novel inhibitory nucleo-cortical circuit controls cerebellar Golgi cell activity. eLife 4

4. Apostolides PF, Trussell LO. 2013. Rapid, activity-independent turnover of vesicular transmitter content at a mixed glycine/GABA synapse. The Journal of neuroscience : the official journal of the Society for Neuroscience 33: 4768–81

5. Aubrey KR, Rossi FM, Ruivo R, Alboni S, Bellenchi GC, et al. 2007. The transporters GlyT2 and VIAAT cooperate to determine the vesicular glycinergic phenotype. The Journal of neuroscience : the official journal of the Society for Neuroscience 27: 6273–81

6. Awatramani GB, Turecek R, Trussell LO. 2005. Staggered development of GABAergic and glycinergic transmission in the MNTB. Journal of neurophysiology 93: 819–28

7. Baade C, Byczkowicz N, Hallermann S. 2016. NMDA receptors amplify mossy fiber synaptic inputs at frequencies up to at least 750 Hz in cerebellar granule cells. Synapse 70: 269–76

8. Bardy C, van den Hurk M, Eames T, Marchand C, Hernandez RV, et al. 2015. Neuronal medium that supports basic synaptic functions and activity of human neurons in vitro. Proceedings of the National Academy of Sciences of the United States of America 112: E2725–34

9. Batten TF, Pow DV, Saha S. 2010. Co-localisation of markers for glycinergic and GABAergic neurones in rat nucleus of the solitary tract: implications for co-transmission. Journal of chemical neuroanatomy 40: 160–76

10. Brandalise F, Gerber U, Rossi P. 2012. Golgi cell-mediated activation of postsynaptic GABA(B) receptors induces disinhibition of the Golgi cell-granule cell synapse in rat cerebellum. PloS one 7: e43417

11. Brickley SG, Cull-Candy SG, Farrant M. 1996. Development of a tonic form of synaptic inhibition in rat cerebellar granule cells resulting from persistent activation of GABAA receptors. The Journal of physiology 497 (Pt 3): 753–9

12. Bright DP, Renzi M, Bartram J, McGee TP, MacKenzie G, et al. 2011. Profound desensitization by ambient GABA limits activation of delta-containing GABAA receptors during spillover. The Journal of neuroscience : the official journal of the Society for Neuroscience 31: 753–63

13. Brown A, Carlyle I, Clark J, Hamilton W, Gibson S, et al. 2001. Discovery and SAR of org 24598-a selective glycine uptake inhibitor. Bioorganic & medicinal chemistry letters 11: 2007–9

14. Cathala L, Misra C, Cull-Candy S. 2000. Developmental profile of the changing properties of NMDA receptors at cerebellar mossy fiber-granule cell synapses. The Journal of neuroscience : the official journal of the Society for Neuroscience 20: 5899–905

15. Caulfield WL, Collie IT, Dickins RS, Epemolu O, McGuire R, et al. 2001. The first potent and selective inhibitors of the glycine transporter type 2. Journal of medicinal chemistry 44: 2679–82

16. Chery N, de Koninck Y. 1999. Junctional versus extrajunctional glycine and GABA(A) receptor-mediated IPSCs in identified lamina I neurons of the adult rat spinal cord. The Journal of neuroscience : the official journal of the Society for Neuroscience 19: 7342–55

17. Chery N, De Koninck Y. 2000. GABA(B) receptors are the first target of released GABA at lamina I inhibitory synapses in the adult rat spinal cord. Journal of neurophysiology 84: 1006–11

18. Crowley JJ, Fioravante D, Regehr WG. 2009. Dynamics of fast and slow inhibition from cerebellar golgi cells allow flexible control of synaptic integration. Neuron 63: 843–53

19. D’Angelo E, Solinas S, Mapelli J, Gandolfi D, Mapelli L, Prestori F. 2013. The cerebellar Golgi cell and spatiotemporal organization of granular layer activity. Frontiers in neural circuits 7: 93

20. Diana MA, Marty A. 2003. Characterization of depolarization-induced suppression of inhibition using paired interneuron--Purkinje cell recordings. The Journal of neuroscience : the official journal of the Society for Neuroscience 23: 5906–18

21. Dieudonne S. 1998. Submillisecond kinetics and low efficacy of parallel fibre-Golgi cell synaptic currents in the rat cerebellum. The Journal of physiology 510 (Pt 3): 845–66

22. DiGregorio DA, Nusser Z, Silver RA. 2002. Spillover of glutamate onto synaptic AMPA receptors enhances fast transmission at a cerebellar synapse. Neuron 35: 521–33

23. DiGregorio DA, Rothman JS, Nielsen TA, Silver RA. 2007. Desensitization properties of AMPA receptors at the cerebellar mossy fiber granule cell synapse. The Journal of neuroscience : the official journal of the Society for Neuroscience 27: 8344–57

24. Dufour A, Tell F, Kessler JP, Baude A. 2010. Mixed GABA-glycine synapses delineate a specific topography in the nucleus tractus solitarii of adult rat. The Journal of physiology 588: 1097–115

25. Dugue GP, Brunel N, Hakim V, Schwartz E, Chat M, et al. 2009. Electrical coupling mediates tunable low-frequency oscillations and resonance in the cerebellar Golgi cell network. Neuron 61: 126–39

26. Dugue GP, Dumoulin A, Triller A, Dieudonne S. 2005. Target-dependent use of co-released inhibitory transmitters at central synapses. The Journal of neuroscience : the official journal of the Society for Neuroscience 25: 6490–8

27. Duguid I, Branco T, Chadderton P, Arlt C, Powell K, Hausser M. 2015. Control of cerebellar granule cell output by sensory-evoked Golgi cell inhibition. Proceedings of the National Academy of Sciences of the United States of America 112: 13099–104

28. Dumba JS, Irish PS, Anderson NL, Westrum LE. 1998. Electron microscopic analysis of gamma-aminobutyric acid and glycine colocalization in rat trigeminal subnucleus caudalis. Brain research 806: 16–25

29. Dumoulin A, Triller A, Dieudonne S. 2001. IPSC kinetics at identified GABAergic and mixed GABAergic and glycinergic synapses onto cerebellar Golgi cells. The Journal of neuroscience : the official journal of the Society for Neuroscience 21: 6045–57

30. Farrant M, Feldmeyer D, Takahashi T, Cull-Candy SG. 1994. NMDA-receptor channel diversity in the developing cerebellum. Nature 368: 335–9

31. Farsi Z, Preobraschenski J, van den Bogaart G, Riedel D, Jahn R, Woehler A. 2016. Single-vesicle imaging reveals different transport mechanisms between glutamatergic and GABAergic vesicles. Science 351: 981–4

32. Fleming E, Hull C. 2019. Serotonin regulates dynamics of cerebellar granule cell activity by modulating tonic inhibition. Journal of neurophysiology 121: 105–14

33. Galliano E, Mazzarello P, D’Angelo E. 2010. Discovery and rediscoveries of Golgi cells. The Journal of physiology 588: 3639–55

34. Gao BX, Stricker C, Ziskind-Conhaim L. 2001. Transition from GABAergic to glycinergic synaptic transmission in newly formed spinal networks. Journal of neurophysiology 86: 492–502

35. Giber K, Diana MA, Plattner V, Dugue GP, Bokor H, et al. 2015. A subcortical inhibitory signal for behavioral arrest in the thalamus. Nature neuroscience 18: 562–68

36. Gonzalez-Forero D, Alvarez FJ. 2005. Differential postnatal maturation of GABAA, glycine receptor, and mixed synaptic currents in Renshaw cells and ventral spinal interneurons. The Journal of neuroscience : the official journal of the Society for Neuroscience 25: 2010–23

37. Granger AJ, Wallace ML, Sabatini BL. 2017. Multi-transmitter neurons in the mammalian central nervous system. Current opinion in neurobiology 45: 85–91

38. Hirono M, Saitow F, Kudo M, Suzuki H, Yanagawa Y, et al. 2012. Cerebellar globular cells receive monoaminergic excitation and monosynaptic inhibition from Purkinje cells. PloS one 7: e29663

39. Hobert O, Glenwinkel L, White J. 2016. Revisiting Neuronal Cell Type Classification in Caenorhabditis elegans. Current biology : CB 26: R1197–R203

40. Holtzman T, Mostofi A, Phuah CL, Edgley SA. 2006a. Cerebellar Golgi cells in the rat receive multimodal convergent peripheral inputs via the lateral funiculus of the spinal cord. The Journal of physiology 577: 69–80

41. Holtzman T, Rajapaksa T, Mostofi A, Edgley SA. 2006b. Different responses of rat cerebellar Purkinje cells and Golgi cells evoked by widespread convergent sensory inputs. The Journal of physiology 574: 491–507

42. Holtzman T, Sivam V, Zhao T, Frey O, van der Wal PD, et al. 2011. Multiple extra-synaptic spillover mechanisms regulate prolonged activity in cerebellar Golgi cell-granule cell loops. The Journal of physiology 589: 3837–54

43. Husson Z, Rousseau CV, Broll I, Zeilhofer HU, Dieudonne S. 2014. Differential GABAergic and glycinergic inputs of inhibitory interneurons and Purkinje cells to principal cells of the cerebellar nuclei. The Journal of neuroscience : the official journal of the Society for Neuroscience 34: 9418–31

44. Ishibashi H, Yamaguchi J, Nakahata Y, Nabekura J. 2013. Dynamic regulation of glycine-GABA co-transmission at spinal inhibitory synapses by neuronal glutamate transporter. The Journal of physiology 591: 3821–32

45. Johnson JW, Ascher P. 1987. Glycine potentiates the NMDA response in cultured mouse brain neurons. Nature 325: 529–31

46. Jonas P, Bischofberger J, Sandkuhler J. 1998. Corelease of two fast neurotransmitters at a central synapse. Science 281: 419–24

47. Kaneda M, Farrant M, Cull-Candy SG. 1995. Whole-cell and single-channel currents activated by GABA and glycine in granule cells of the rat cerebellum. The Journal of physiology 485 (Pt 2): 419–35

48. Keller AF, Coull JA, Chery N, Poisbeau P, De Koninck Y. 2001. Region-specific developmental specialization of GABA-glycine cosynapses in laminas I-II of the rat spinal dorsal horn. The Journal of neuroscience : the official journal of the Society for Neuroscience 21: 7871–80

49. Kepecs A, Fishell G. 2014. Interneuron cell types are fit to function. Nature 505: 318–26

50. Koga R, Miyoshi Y, Sakaue H, Hamase K, Konno R. 2017. Mouse d-Amino-Acid Oxidase: Distribution and Physiological Substrates. Frontiers in molecular biosciences 4: 82

51. Kozareva V, Martin C, Osorno T, Rudolph S, Guo C, et al. 2021. A transcriptomic atlas of mouse cerebellar cortex comprehensively defines cell types. Nature 598: 214–19

52. Lim L, Mi D, Llorca A, Marin O. 2018. Development and Functional Diversification of Cortical Interneurons. Neuron 100: 294–313

53. Lu T, Rubio ME, Trussell LO. 2008. Glycinergic transmission shaped by the corelease of GABA in a mammalian auditory synapse. Neuron 57: 524–35

54. Mapelli L, Rossi P, Nieus T, D’Angelo E. 2009. Tonic activation of GABAB receptors reduces release probability at inhibitory connections in the cerebellar glomerulus. Journal of neurophysiology 101: 3089–99

55. Mapelli L, Solinas S, D’Angelo E. 2014. Integration and regulation of glomerular inhibition in the cerebellar granular layer circuit. Frontiers in cellular neuroscience 8: 55

56. McIntire SL, Reimer RJ, Schuske K, Edwards RH, Jorgensen EM. 1997. Identification and characterization of the vesicular GABA transporter. Nature 389: 870–6

57. Mitchell SJ, Silver RA. 2000. GABA spillover from single inhibitory axons suppresses low-frequency excitatory transmission at the cerebellar glomerulus. The Journal of neuroscience : the official journal of the Society for Neuroscience 20: 8651–8

58. Mitchell SJ, Silver RA. 2003. Shunting inhibition modulates neuronal gain during synaptic excitation. Neuron 38: 433–45

59. Monyer H, Burnashev N, Laurie DJ, Sakmann B, Seeburg PH. 1994. Developmental and regional expression in the rat brain and functional properties of four NMDA receptors. Neuron 12: 529–40

60. Moore LA, Trussell LO. 2017. Corelease of Inhibitory Neurotransmitters in the Mouse Auditory Midbrain. The Journal of neuroscience : the official journal of the Society for Neuroscience 37: 9453–64

61. Nabekura J, Katsurabayashi S, Kakazu Y, Shibata S, Matsubara A, et al. 2004. Developmental switch from GABA to glycine release in single central synaptic terminals. Nature neuroscience 7: 17–23

62. Negyessy L, Vidnyanszky Z, Kuhn R, Knopfel T, Gorcs TJ, Hamori J. 1997. Light and electron microscopic demonstration of mGluR5 metabotropic glutamate receptor immunoreactive neuronal elements in the rat cerebellar cortex. The Journal of comparative neurology 385: 641–50

63. Neki A, Ohishi H, Kaneko T, Shigemoto R, Nakanishi S, Mizuno N. 1996. Metabotropic glutamate receptors mGluR2 and mGluR5 are expressed in two non-overlapping populations of Golgi cells in the rat cerebellum. Neuroscience 75: 815–26

64. Nerlich J, Rubsamen R, Milenkovic I. 2017. Developmental Shift of Inhibitory Transmitter Content at a Central Auditory Synapse. Frontiers in cellular neuroscience 11: 211

65. Nietz AK, Vaden JH, Coddington LT, Overstreet-Wadiche L, Wadiche JI. 2017. Non-synaptic signaling from cerebellar climbing fibers modulates Golgi cell activity. eLife 6

66. Nusbaum MP, Blitz DM, Marder E. 2017. Functional consequences of neuropeptide and small-molecule co-transmission. Nature reviews. Neuroscience 18: 389–403

67. O’Brien JA, Berger AJ. 1999. Cotransmission of GABA and glycine to brain stem motoneurons. Journal of neurophysiology 82: 1638–41

68. Ottersen OP, Davanger S, Storm-Mathisen J. 1987. Glycine-like immunoreactivity in the cerebellum of rat and Senegalese baboon, Papio papio: a comparison with the distribution of GABA-like immunoreactivity and with [3H]glycine and [3H]GABA uptake. Experimental brain research 66: 211–21

69. Ottersen OP, Storm-Mathisen J, Somogyi P. 1988. Colocalization of glycine-like and GABA-like immunoreactivities in Golgi cell terminals in the rat cerebellum: a postembedding light and electron microscopic study. Brain research 450: 342–53

70. Paik SK, Yoo HI, Choi SK, Bae JY, Park SK, Bae YC. 2019. Distribution of excitatory and inhibitory axon terminals on the rat hypoglossal motoneurons. Brain structure & function

71. Parra P, Gulyas AI, Miles R. 1998. How many subtypes of inhibitory cells in the hippocampus? Neuron 20: 983–93

72. Petilla Interneuron Nomenclature G, Ascoli GA, Alonso-Nanclares L, Anderson SA, Barrionuevo G, et al. 2008. Petilla terminology: nomenclature of features of GABAergic interneurons of the cerebral cortex. Nature reviews. Neuroscience 9: 557–68

73. Powell K, Mathy A, Duguid I, Hausser M. 2015. Synaptic representation of locomotion in single cerebellar granule cells. eLife 4

74. Riquelme R, Saldana E, Osen KK, Ottersen OP, Merchan MA. 2001. Colocalization of GABA and glycine in the ventral nucleus of the lateral lemniscus in rat: an in situ hybridization and semiquantitative immunocytochemical study. The Journal of comparative neurology 432: 409–24

75. Rossi DJ, Hamann M. 1998. Spillover-mediated transmission at inhibitory synapses promoted by high affinity alpha6 subunit GABA(A) receptors and glomerular geometry. Neuron 20: 783–95

76. Rousseau CV, Dugue GP, Dumoulin A, Mugnaini E, Dieudonne S, Diana MA. 2012. Mixed inhibitory synaptic balance correlates with glutamatergic synaptic phenotype in cerebellar unipolar brush cells. The Journal of neuroscience : the official journal of the Society for Neuroscience 32: 4632–44

77. Rousseau F, Aubrey KR, Supplisson S. 2008. The glycine transporter GlyT2 controls the dynamics of synaptic vesicle refilling in inhibitory spinal cord neurons. The Journal of neuroscience : the official journal of the Society for Neuroscience 28: 9755–68

78. Saunders A, Macosko EZ, Wysoker A, Goldman M, Krienen FM, et al. 2018. Molecular Diversity and Specializations among the Cells of the Adult Mouse Brain. Cell 174: 1015–30 e16

79. Scala F, Kobak D, Bernabucci M, Bernaerts Y, Cadwell CR, et al. 2020. Phenotypic variation of transcriptomic cell types in mouse motor cortex. Nature

80. Schonewille M, Girasole AE, Rostaing P, Mailhes-Hamon C, Ayon A, et al. 2021. NMDARs in granule cells contribute to parallel fiber-Purkinje cell synaptic plasticity and motor learning. Proceedings of the National Academy of Sciences of the United States of America 118

81. Schwartz EJ, Rothman JS, Dugue GP, Diana M, Rousseau C, et al. 2012. NMDA receptors with incomplete Mg(2)(+) block enable low-frequency transmission through the cerebellar cortex. The Journal of neuroscience : the official journal of the Society for Neuroscience 32: 6878–93

82. Silver RA, Traynelis SF, Cull-Candy SG. 1992. Rapid-time-course miniature and evoked excitatory currents at cerebellar synapses in situ. Nature 355: 163–6

83. Simat M, Parpan F, Fritschy JM. 2007. Heterogeneity of glycinergic and gabaergic interneurons in the granule cell layer of mouse cerebellum. The Journal of comparative neurology 500: 71–83

84. Simpson JI, Hulscher HC, Sabel-Goedknegt E, Ruigrok TJ. 2005. Between in and out: linking morphology and physiology of cerebellar cortical interneurons. Progress in brain research 148: 329–40

85. Stell BM, Brickley SG, Tang CY, Farrant M, Mody I. 2003. Neuroactive steroids reduce neuronal excitability by selectively enhancing tonic inhibition mediated by delta subunit-containing GABAA receptors. Proceedings of the National Academy of Sciences of the United States of America 100: 14439–44

86. Supplisson S, Bergman C. 1997. Control of NMDA receptor activation by a glycine transporter co-expressed in Xenopus oocytes. The Journal of neuroscience : the official journal of the Society for Neuroscience 17: 4580–90

87. Szoboszlay M, Lorincz A, Lanore F, Vervaeke K, Silver RA, Nusser Z. 2016. Functional Properties of Dendritic Gap Junctions in Cerebellar Golgi Cells. Neuron 90: 1043–56

88. Tasic B, Menon V, Nguyen TN, Kim TK, Jarsky T, et al. 2016. Adult mouse cortical cell taxonomy revealed by single cell transcriptomics. Nature neuroscience 19: 335–46

89. Todd AJ, Sullivan AC. 1990. Light microscope study of the coexistence of GABA-like and glycine-like immunoreactivities in the spinal cord of the rat. The Journal of comparative neurology 296: 496–505

90. Tremblay R, Lee S, Rudy B. 2016. GABAergic Interneurons in the Neocortex: From Cellular Properties to Circuits. Neuron 91: 260–92

91. Vervaeke K, Lorincz A, Gleeson P, Farinella M, Nusser Z, Silver RA. 2010. Rapid desynchronization of an electrically coupled interneuron network with sparse excitatory synaptic input. Neuron 67: 435–51

92. Virtanen P, Gommers R, Oliphant TE, Haberland M, Reddy T, et al. 2020. SciPy 1.0: fundamental algorithms for scientific computing in Python. Nature methods 17: 261–72

93. Vos BP, Volny-Luraghi A, De Schutter E. 1999. Cerebellar Golgi cells in the rat: receptive fields and timing of responses to facial stimulation. The European journal of neuroscience 11: 2621–34

94. Wall MJ, Usowicz MM. 1997. Development of action potential-dependent and independent spontaneous GABAA receptor-mediated currents in granule cells of postnatal rat cerebellum. The European journal of neuroscience 9: 533–48

95. Wang L, Tu P, Bonet L, Aubrey KR, Supplisson S. 2013. Cytosolic transmitter concentration regulates vesicle cycling at hippocampal GABAergic terminals. Neuron 80: 143–58

96. Wang LZ, Zhu XZ. 2003. Spatiotemporal relationships among D-serine, serine racemase, and D-amino acid oxidase during mouse postnatal development. Acta pharmacologica Sinica 24: 965–74

97. Watanabe D, Nakanishi S. 2003. mGluR2 postsynaptically senses granule cell inputs at Golgi cell synapses. Neuron 39: 821–9

98. Wolosker H, Blackshaw S, Snyder SH. 1999. Serine racemase: a glial enzyme synthesizing D-serine to regulate glutamate-N-methyl-D-aspartate neurotransmission. Proceedings of the National Academy of Sciences of the United States of America 96: 13409–14

99. Yuste R, Hawrylycz M, Aalling N, Aguilar-Valles A, Arendt D, et al. 2020. A community-based transcriptomics classification and nomenclature of neocortical cell types. Nature neuroscience 23: 1456–68

100. Zafra F, Aragon C, Olivares L, Danbolt NC, Gimenez C, Storm-Mathisen J. 1995a. Glycine transporters are differentially expressed among CNS cells. The Journal of neuroscience : the official journal of the Society for Neuroscience 15: 3952–69

101. Zafra F, Gomeza J, Olivares L, Aragon C, Gimenez C. 1995b. Regional distribution and developmental variation of the glycine transporters GLYT1 and GLYT2 in the rat CNS. The European journal of neuroscience 7: 1342–52

102. Zeilhofer HU, Studler B, Arabadzisz D, Schweizer C, Ahmadi S, et al. 2005. Glycinergic neurons expressing enhanced green fluorescent protein in bacterial artificial chromosome transgenic mice. The Journal of comparative neurology 482: 123–41

103. Zeisel A, Munoz-Manchado AB, Codeluppi S, Lonnerberg P, La Manno G, et al. 2015. Brain structure. Cell types in the mouse cortex and hippocampus revealed by single-cell RNA-seq. Science 347: 1138–42

104. Zeng H, Sanes JR. 2017. Neuronal cell-type classification: challenges, opportunities and the path forward. Nature reviews. Neuroscience 18: 530–46

